# Characterization of trade-offs between immunity and reproduction in *Astrangia poculata*

**DOI:** 10.1101/2023.07.24.550370

**Authors:** Natalie Villafranca, Isabella Changsut, Sofia C. Diaz de Villegas, Haley R. Womack, Lauren E. Fuess

## Abstract

**Background:** Living organisms face ubiquitous pathogenic threats and have consequently evolved immune systems to protect against potential invaders. However, many components of the immune system are physiologically costly to maintain and engage, often drawing resources away from other organismal processes such as growth and reproduction. Evidence from a diversity of systems has demonstrated that organisms use complex resource allocation mechanisms to manage competing needs and optimize fitness. However, understanding of resource allocation patterns is limited across taxa. Cnidarians, which include ecologically important organisms like hard corals, have been historically understudied in the context of resource allocations. Improving understanding of resource allocation-associated tradeoffs in cnidarians is critical for understanding future ecological dynamics in the face of rapid environmental change.

**Methods:** Here, we characterize trade-offs between constitutive immunity and reproduction in the facultatively symbiotic coral Astrangia poculata. Male colonies underwent ex situ spawning and sperm output was quantified. We then examined the effects of variable symbiont density and energetic budget on physiological traits, including immune activity and reproductive investment. Furthermore, we tested for potential trade-offs between immune activity and reproductive investment.

**Results:** We found limited effects of energetic budget on immune metrics; melanin production was significantly positively associated with energetic budget. However, we failed to document any associations between immunity and reproductive output which would be indicative of trade-offs, possibly due to experimental limitations. Our results provide a preliminary framework for future studies investigating immune trade-offs in cnidarians.

## Introduction

The ability to successfully defend against a variety of pathogenic threats is an essential component of organismal fitness (Lochmiller & Deerenberg 2000). However, most immune defenses which prevent or mitigate damage from pathogens are energetically costly (Lochmiller & Deerenberg 2000). This is particularly important as organisms operate using a fixed budget of energetic resources that must be allocated to competing demands (i.e. growth, reproduction, defense, etc; (Stearns 1989)). Consequently, allocation of resources to one demand often comes at the cost of others, creating trade-offs (Melbinger & Vergassola 2015). Optimal organismal fitness therefore requires careful allocation of resources across these demands to minimize negative effects of tradeoffs and maximize reproductive output. Notably, the high costs of maintaining immune systems and mounting immune responses often result in trade-offs with other traits (Lochmiller & Deerenberg 2000; Martin et al. 2008; Rauw 2012; Rigby & Jokela 2000; van der Most et al. 2010). Among the most well documented immune-related tradeoffs are those between reproduction and immunity (Adamo et al. 2001; Brokordt et al. 2019; Gwynn et al. 2005; Hosken 2001; Schwenke et al. 2016). However, despite frequent empirical observation of immune-reproductive trade-offs and robust proposed theory, many questions still remain regarding the generalizability of resource allocation theory and immune-related tradeoffs. In some systems, consistent immune-reproductive tradeoffs have not been observed (Kelley et al. 2021; McNamara et al. 2013; Miyashita et al. 2019; Syed et al. 2020; Xu et al. 2012). Furthermore, it remains unclear how dynamic changes in resources/energetic budget might impact these trade-offs. It is theorized that since total energy and relative allocation are co-dependent, variation in energy budget may have an impact on trade-offs such as those between immunity and reproduction, however limited empirical evidence exists to support or refute this theory (Cotter et al. 2010; Descamps et al. 2016; Simmons 2012; Stahlschmidt et al. 2013). Combined, these gaps in existing theory make it difficult to predict how resources are allocated among traits, requiring further study to improve understanding of resource allocation and immune trade-offs in complex contexts. This is particularly important in light of the rising prevalence of epizootic outbreaks affecting vulnerable species across the globe (Croquer & Weil 2009; Glenn & Pugh 2006; Kilburn et al. 2010).

Scleractinian or reef building corals are arguably one of the taxa most impacted by increases in epizootic outbreaks (Bruno et al. 2007; Precht et al. 2016; Ruiz-Moreno et al. 2012). Disease outbreaks have been one of the largest drivers of coral declines in recent decades, affecting almost all major reef building species (Alvarez-Filip et al. 2022; Bourne et al. 2009; Cróquer et al. 2021; Moriarty et al. 2020; Sharma & Ravindran 2020; Sutherland et al. 2004). Still, despite the severity of these outbreaks, many gaps remain in our understanding of associated disease processes. For example, significant inter- and intra-specific variation has been observed in coral disease resilience (Alvarez-Filip et al. 2022; Miller et al. 2019; Palmer et al. 2011). While some studies have pointed to the importance of divergence in certain cellular processes in driving this variation (Beavers et al. 2023; Fuess et al. 2017; MacKnight et al. 2022; Pinzon et al. 2014a; Traylor-Knowles et al. 2021), little is known regarding how broader ecological processes, including resource allocation, might contribute to variation in disease resistance. To date, most studies of cnidarian resource allocation have focused on stress associated trade-offs. The negative effects of thermal anomalies on coral reproductive output have been well documented (Michalek-Wagner & Willis 2001; Nielsen et al. 2020; Paxton et al. 2016). In contrast, only a handful of studies have documented reproductive tradeoffs involving immunological processes, all of which had focused on the negative impacts of disease outbreaks on coral reproduction (Fuess et al. 2018; Weil et al. 2009). No studies have directly linked reproductive investment and constitutive immunity in cnidarians, in part due to the historic lack of tractable model cnidarian systems.

Recent advances in the development of cnidarian model and study systems have greatly expanded the scope of cnidarian research. One such emergent model species is *Astrangia poculata* (common name: northern star coral), a non-reef building stony coral that can be found along the Atlantic Coast of the United States ranging from Cape Cod, MA to the Texas Gulf Coast (Dimond et al. 2012). Importantly, unlike tropical corals which are dependent on symbiotic relationships for survival, *A. poculata* associates facultatively with a single species of symbiont, *Breviolum psygmophilum* (Lajeunesse et al. 2012) with colonies ranging from high symbiont density (“brown”) to low symbiont density (“white”; (Sharp et al. 2017). Classification of symbiotic states is based on color, approximate chlorophyll concentration, and symbiont density (“brown” > 10^6^ cells cm^−2^; “white” 10^4^ −10^6^ cells cm^−2^; Sharp et al. 2017). A key benefit of the symbiotic relationship between corals and their symbionts is the exchange of organic nutrients to the host (Kirk N.L. 2016). Consequently, facultative symbiosis may serve as a natural system for exploring the effects of variable resource budget on immune-associated tradeoffs in corals, though the exact effects of variation in symbiont density on host energetic budget are poorly studied (Szmant-Froelich & Pilson 1980).

Here we used the tractable *A. poculata* study system to investigate cnidarian resource allocation and potential trade-offs between reproduction and immunity. We first assessed general effects of variability in symbiont density on host physiology (reproductive output, energetic budget, and immune activity). Next, we assessed the effects of energetic budget on both reproductive output and immune activity generally, independent of symbiont density. Finally, we tested for reproductive-immune tradeoffs. Our study provides a preliminary framework for future studies considering the inter-connected roles of symbiosis, resource allocation, and immunity in cnidarians.

## Materials & Methods

### Sample Collection & Experimental Design

Colonies of “white” (low symbiont density) and “brown” (high symbiont densities) *A. poculata* were collected from Fort Wetherill in Jamestown, Rhode Island (41°28′40″ N, 71°21′34″ W) in August 2021, at a depth of 10-15 meters. Corals were collected under RI Department of Environmental Management Permit #825 issued to Koty Sharp/Roger Williams University. Previous studies have indicated *A. poculata* reaches peak gametogenesis between August and September each year (Szmant-Froelich et al. 1980). Samples were returned to Roger Williams University (Bristol, Rhode Island), and held overnight prior to experimental spawning. To trigger spawning, individuals were moved from the recirculating tanks at ambient temperatures (∼19°C) to individual containers with 100 mL of filtered seawater heated to 27°C. Colonies were then closely monitored for signs of spawning. Colony sex was confirmed based on gametes released at time of spawning (*A. poculata* is gonochoristic (Szmant-Froelich et al. 1980)). Twenty male colonies were identified and used for the study, 13 white and 7 brown. Individuals were allowed to spawn for 30 minutes, at which point gamete release had nearly or completely ceased. Following this period adult colonies were removed and flash frozen in liquid nitrogen. The water in the container was then thoroughly mixed and a 1 mL sample was taken from each colony for sperm density estimation. Sperm density was estimated by counting in triplicate on a haemocytometer; counts were then normalized to number of polyps per colony prior to statistical analyses. Flash frozen adult corals were later shipped to Texas State University for downstream sample processing and analysis.

### Symbiont Density Quantification

Flash frozen adult colonies were airbrushed using a Paasche air brusher (VL0221) with 100 mM Tris + 0.5 mM Dithiothreitol (DTT; pH 7.8) to remove tissue from the skeleton and extract symbionts and host-enriched proteins (Fuess et al. 2016). First, in order to estimate symbiont density, an area of 2.14 cm^2^ (1-2 polyps) was marked on a flat surface of the colony. Tissue was airbrushed from the area until no tissue remained. The resulting tissue slurry was placed in a 2.0 mL microcentrifuge tube, and homogenized for 10 seconds using a handheld homogenizer (Fisherbrand 150). Following homogenization, samples were centrifuged (2000 RPM for 3 minutes) and then washed with 500 uL of DI H_2_O. This process was repeated twice; final samples were preserved in 500 uL of Deionized H_2_O + 0.01% Sodium Dodecyl Sulfate (SDS) and stored at 4°C. Symbiont density was later determined via microscopy (Nikon Eclipse E600) by counting in triplicate on a hemocytometer (VWR Scientific Counting Chamber, cat #15170-208; (Changsut et al. 2022; Mieog et al. 2009)).

### Protein Extraction

Tissues for protein extraction were removed from the remaining portion of the *A. poculata* colony via airbrushing with 100 mM Tris + 0.5 mM DTT (pH 7.8). The resultant slury was homogenized for one minute, followed by a 10 minute incubation on ice. One milliliter of resulting slurry was reserved for melanin concentration estimation, stored at −20°C. The remaining volume was centrifuged for 5 minutes (3500 RPM at 4°C). The resultant supernatant was stored at −80°C for later analyses.

Immune, lipid, and carbohydrate assays were performed using preserved host-enriched protein and melanin aliquots. Colorimetric immune assays were run in triplicate on 96 well plates using a Cytation 1 cell imaging multi-mode reader and Gen 5 software (BioTek Instruments, Winooski, VT, USA). A Red660 (G Biosciences, St. Louis, Missouri) assay was used to determine protein concentration to standardize immune activity metrics (Mydlarz & Palmer 2011).

### Prophenoloxidase cascade assays (Total PO and Melanin)

Total phenoloxidase activity (prophenoloxidase + phenoloxidase) was measured by diluting 20 uL of coral protein extract in 20 uL of 50 mM PBS (pH 7.0) in a 96 well plate (Costar, Corning, Kennebunk, ME). In order to convert prophenoloxidase to phenoloxidase, wells were then incubated with 25 uL of trypsin (0.1 mg/mL DI) for 30 minutes at room temperature. Negative controls were run in duplicate using extraction buffer (100 mM Tris + 0.5 mM DTT (pH 7.8)) in place of sample. Following incubation, 30 uL of dopamine (10 mM) was added to each sample to initiate the reaction. Immediately after addition of dopamine, absorbance was read every minute for 20 minutes at 490 nm. Total PO activity was calculated as change of absorbance (final – initial) per milligrams protein per minute, at the steepest point of the curve, standardized by protein concentration (Mydlarz & Palmer 2011).

Tissue samples for melanin analysis were desiccated in a SpeedVac (Eppendorf, Vacufuge plus). To calculate melanin concentration, a spatula full of 10 mm glass beads (∼200 uL) was added to each tube, and vortexed for 10 seconds. Next, 400 uL of 10M NaOH were added, and samples were vortexed again for 20 seconds. Samples then underwent a 48 hour incubation period, with vortexing at 24 and 48 hours. At the end of this period, tubes were centrifuged at 1000 RPM for 10 minutes at room temperature. Forty microliters of the supernatant were transferred to a ½ well 96-well UV plate (UV-STAR, Greiner bio-one, Frickenhausen, Germany). A standard curve of melanin dissolved in 10 M NaOH and processed identically to the samples was used to convert absorbance readings to melanin concentration (Mydlarz & Palmer 2011). Absorbance was recorded at 410 and 490 nm. Melanin concentrations were calculated using a melanin standard (Sigma Aldrich, St. Louis, Missouri) and standardized to total dry tissue weight (mg melanin/mg tissue).

### Antioxidant assays

Following previously established protocols, activity was assessed for two antioxidants: catalase (CAT) and peroxidase (POX; (Fuess et al. 2016; Mydlarz & Palmer 2011)). Catalase activity was measured by adding 5 uL of sample protein extract to a 96-well UV well plate (UV-STAR, Greiner bio-one, Frickenhausen, Germany), which was diluted with 45 uL of 50 mM PBS (pH 7.0). Seventy-five microliters of 25 mM H_2_O_2_ were added to each well to initiate the reaction. Absorbance was immediately measured at 240 nm every 30 seconds for 15 minutes. A standard curve of hydrogen peroxide was used to calculate H_2_O_2_ concentrations. Catalase activity was calculated as H_2_O_2_ scavenged per min at the steepest point of the curve, normalized to protein concentration (Fuess et al. 2016; Mydlarz & Palmer 2011).

Peroxidase activity was measured by diluting 20 uL of sample protein extract with 20 uL of 10 mM PBS (pH 6.0) in a 96-well plate (Costar, Corning, Kennebunk, ME). Then, 25 uL of 5 mM guaiacol in 10 mM PBS (pH 6.0) was added. Negative controls were run in duplicate using extraction buffer (100 mM Tris + 0.5 mM DTT (pH 7.8)) in place of protein extract. To initiate the reaction, 20 uL of 20 mM H_2_O_2_ was pipetted into each well. Absorbance was read at 470 nm every minute for 15 minutes. Activity was calculated as change in absorbance per minute at the steepest point of the curve, normalized to protein concentration (Mydlarz & Harvell 2007).

### Antibacterial activity

Antibacterial activity was assessed against *Vibrio coralliilyticus* (Strain RE22Sm provided by D. Nelson at University of Rhode Island) grown in Luria Broth. *V. coralliilyticus* is a known cnidarian pathogen with roles in numerous coral disease (Ushijima et al. 2020; Ushijima et al. 2014). In a sterile hood, 140 uL of *V. coralliilyticus* diluted to a optical density of 0.2 at 600 nm and 60 uL of sample diluted to a standard protein concentration were added to wells of a sterile 96-well plate. The plate was incubated for 6 hours at 27°C, with absorbance at 600 nm recorded every 10 minutes. Bacterial growth rates were determined as the change in absorbance during the logarithmic growth phase (Pinzon et al. 2014b).

### Total Lipid Content

A standard protocol for quantification of lipids within coral tissue slurries was used to estimate lipid content (Bove & Baumann 2021). Aliquots of 150 uL of sample extract were desiccated overnight in a SpeedVac (Eppendorf, Vacufuge plus). Five hundred microliters of a 2:1 chloroform/methanol mixture and 100 uL of 0.05 M NaCl were added to each desiccated sample. Samples were vortexed to dissolve lipids, and were placed on a shaker (Corning LSE Benchtop Shaking Incubator) for one hour, vortexing every 15 minutes. Samples were then centrifuged at 3000 RPM for 5 minutes. One hundred microliters of the bottom layer of the centrifuged samples were then added to a 96-well PCR plate in triplicate. Fifty microliters of methanol were added to all wells and placed in a hot water bath (70°C) for 15 minutes. Following solvent evaporation, 100 uL of 18 M sulfuric acid were added to each well, and placed in a thermocycler (Eppendorf Mastercycler Nexus) at 90°C for 20 minutes followed by 4°C for 20 minutes. Seventy-five microliters of sample were transferred from each PCR well into a new 96 well plate. Following an initial absorbance reading at 540 nm, 34.5 uL of 0.2 mg/mL vanillin in 17% phosphoric acid were added to each well, and incubated in the dark at room temperature for 5 minutes, after which absorbance was read again (Cheng et al. 2011). The difference between the resulting absorbance was converted to lipid concentration (mg/mL) using a standard curve of corn oil diluted in chloroform treated identically to samples. Final values were standardized based on dry tissue weight (lipids/ug tissue).

### Total Carbohydrate Content

Total carbohydrate concentration was estimated following previously established protocols (Dubois et al. 1956; Masuko et al. 2005). Fifty microliters of sample extract, blank, or glucose standard were added in triplicate to a 96 well plate. 150 uL of H_2_SO_4_ were added to each well, immediately followed by 30 uL of 5% phenol. The plate was incubated in an uncovered hot water bath (70°C) for 5 minutes, and cooled for 15 minutes. Absorbance was read at 485 nm and 750 nm, and then the absorbance values at 750 nm were subtracted from the absorbance values at 485 nm to correct for differential scattering (Fu et al. 2008). A serial dilution of glucose, treated identically to samples, was used to convert absorbance to total carbohydrate concentration.

### Statistical Analyses

All statistical analyses were performed in RStudio (2021.09.1). Prior to all statistical tests, outliers were removed (n=1 each for peroxidase, melanin, and lipid assay). Data was checked for normality, and total phenoloxidase activity was normalized using a log transformation. First we considered the effects of symbiont density on sperm output, energetic budget, and immune metric activity using linear models. To test for the effects of symbiont density on sperm output we ran the following linear model: sperm output ∼ symbiont density, where symbiont density is the square root normalized amount of symbionts per area (cm^2^) and sperm output is the average sperm output per polyp. Then we ran the linear models: lipid concentration ∼ symbiont density and carbohydrate concentration ∼ symbiont density; total lipid and total carbohydrate were reflective of concentration standardized by dry tissue weight. Finally, we ran the linear model: immune metric ∼ symbiont density for each measured immune metric individual, using standardized immune activity assay as the response variable.

Next, we tested the effects of energetic budget on both reproductive output and immunity. To start, we assessed impacts of energetic budget on sperm output using the linear models: sperm output ∼ total lipid and sperm output ∼ total carbohydrate. We assessed the effects of variation in energetic budget on immune metric activity using the models: immune activity ∼ lipid concentration and immune activity ∼ carbohydrate activity. For immune assays which were significantly associated with energetic budget, we also tested for effects of symbiont state (white or brown) on this relationship using the linear models: immune activity ∼ lipid concentration*type and immune activity ∼ carbohydrate concentration*type. Linear regressions were plotted using the R package ggplot2 (Wickham 2016). Finally, we tested for potential immune-reproduction tradeoffs using individual linear models for each immune metric: immune activity ∼ sperm output.

## Results

We first considered impacts of symbiont density on lipid and carbohydrate concentrations as well as immune metric activity. We found no significant association between symbiont density and lipid concentration (linear model, est = −1.17e^-6^, std err = 3.42E^-6^, *p* = 0.736) or carbohydrate concentration (linear model, est = −2.13e^-6^, std err = 4.28e^-6^, *p* = 0.626). Furthermore, symbiont density did not have significant effects on sperm output (linear model, est. = 5.69e^-4^, std err= 4.64e^-3^, *p* = 0.901). Finally, we considered associations between symbiont density and activity of our measured immune metrics; symbiont density did not significantly impact activity of any of our immune metrics (**Table 1**).

**Table 1:** Result of linear model of the effects of symbiont density on measured immune metrics.

| Response Variable | estimate | std err | t value | p value |
| --- | --- | --- | --- | --- |
| Catalase | 0.275 | 0.342 | 0.804 | 0.432 |
| Peroxidase | -1.24e<sup>-3</sup> | 1.22e<sup>-3</sup> | -1.02 | 0.324 |
| Total Phenoloxidase | -2.10e<sup>-4</sup> | 6.05e<sup>-4</sup> | -0.348 | 0.732 |
| Melanin | 7.34e<sup>-8</sup> | 4.19e<sup>-8</sup> | 1.75 | 0.0975 |
| Antibacterial Activity | -2.80e<sup>-4</sup> | 3.72e<sup>-4</sup> | -0.754 | 0.461 |

Next, we considered the effects of energetic budget (lipid and carbohydrate concentration) on both sperm output and immune assay activity. Sperm output was not associated with either lipid concentration (linear model, est = 401.58, std err = 325.38, *p* = 0.221), or carbohydrate concentration (linear model, est = 423.88, std err = 233.82, *p* = 0866). Melanin concentration was significantly positively associated with both lipid and carbohydrate concentration (**Fig 1**, **Tables 2-3**). No other immune metric was associated with either of our metrics of energetic budget (**Tables 2-3**). Furthermore, the association between melanin and lipid concentration as well as between melanin and carbohydrate concentration did not vary as a result of symbiotic state (**Tables 4-5**; white versus brown). Finally, we assessed potential trade-offs between reproductive output and immunity. None of our measured immune metrics were significantly associated with sperm output (**Table 6**).

**Fig 1:**
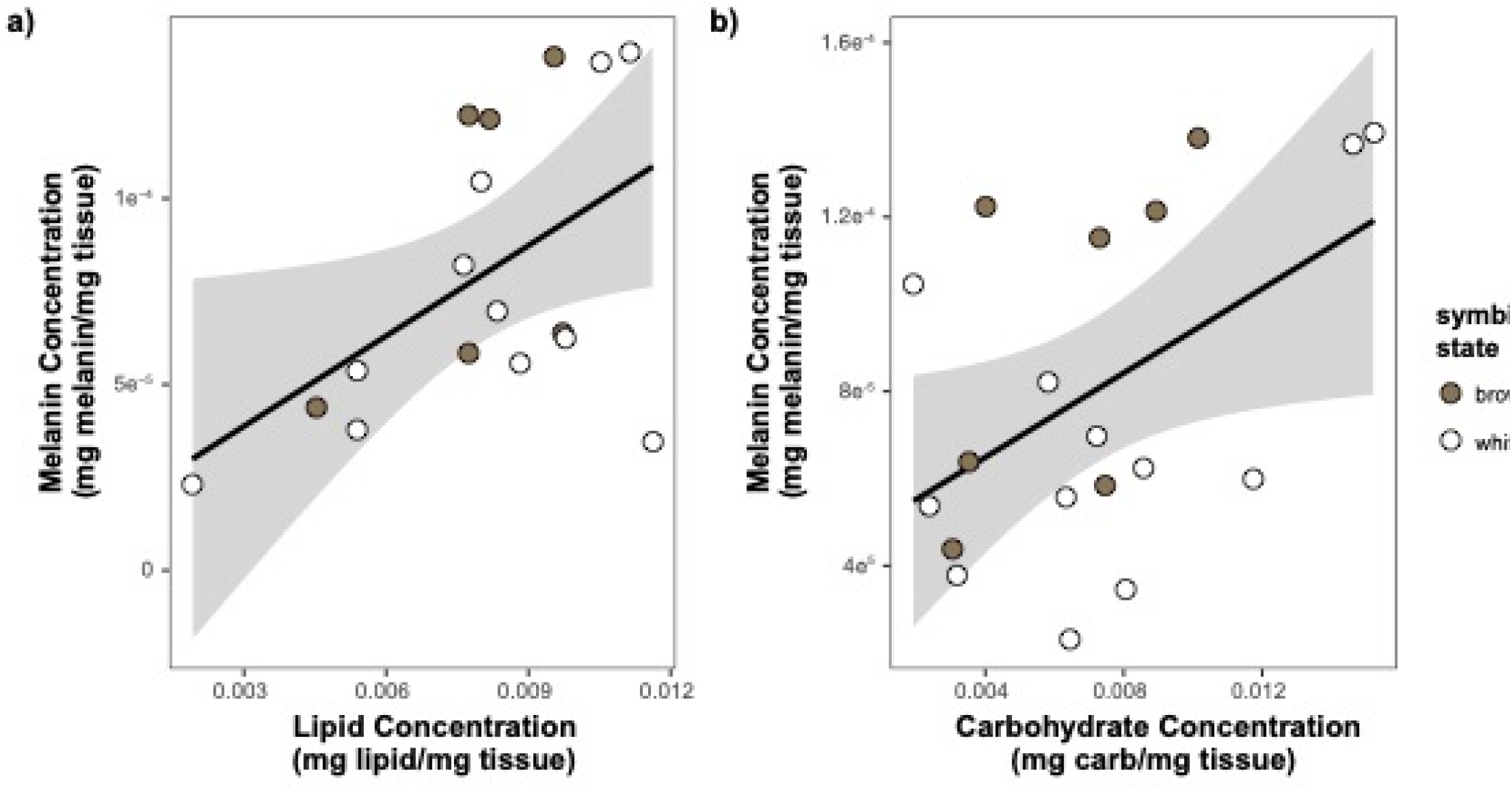
results of linear regression modeling of the relationship between **a)** lipid and **b)** carbohydrate concentration and melanin concentration. Points are colored based on original sample symbiotic state classifcaiton (white or brown). Trendline is representative of the linear model of the relationship of the two variables, with 95% confidence intervals shaded.

**Table 2:** Results of linear model of the effects of lipid concentration on measured immune metrics.

| Response Variable | estimate | std err | t value | p value |
| --- | --- | --- | --- | --- |
| Catalase | -3948 | 31105 | -0.127 | 0.901 |
| Peroxidase | 37.3 | 112.6 | 0.331 | 0.745 |
| Total Phenoloxidase | 49.6 | 51.2 | 0.970 | 0.346 |
| Melanin | -8.10e<sup>-3</sup> | 3.47e<sup>-3</sup> | 2.34 | 0.0338* |
| Antibacterial Activity | 14.8 | 31.9 | 0.463 | 0.65 |

**Table 3:**
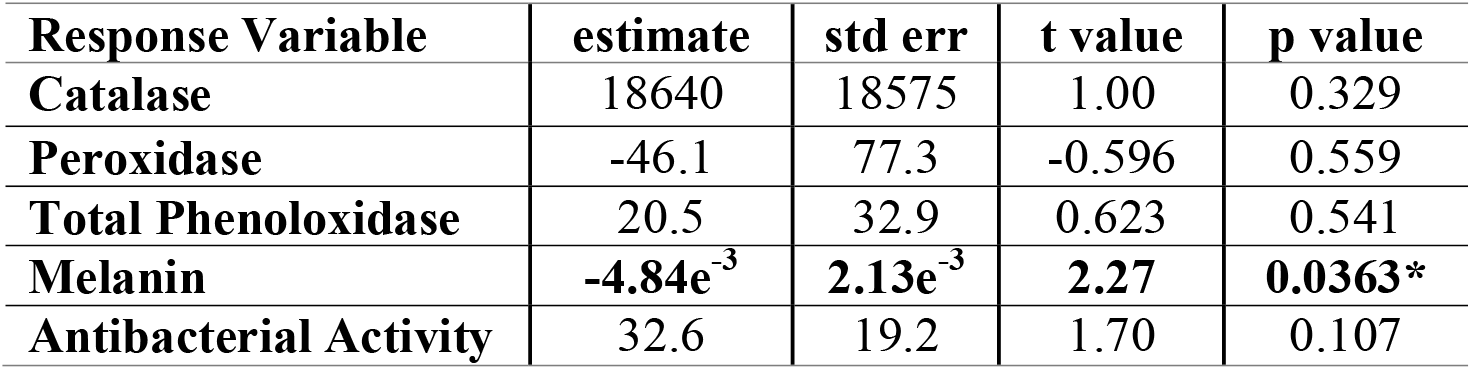
Results of linear model of the effects of carbohydrate concentration on measured immune metrics.

**Table 4:** Results of linear model of the interactive effects of lipid concentration and symbiont state on melanin concentration.

| Predictor | estimate | std err | t value | p value |
| --- | --- | --- | --- | --- |
| Lipid | 1.12e<sup>-2</sup> | 8.55e<sup>-3</sup> | 1.31 | 0.214 |
| Type | 8.64e<sup>-6</sup> | 7.68e<sup>-5</sup> | 0.113 | 0.912 |
| Lipid*Type | -3.58e<sup>-3</sup> | 9.40e<sup>-3</sup> | -0.381 | 0.709 |

**Table 5:** Results of linear model of the interactive effects of carbohydrate concentration and symbiont state on melanin concentration.

| Predictor | estimate | std err | t value | p value |
| --- | --- | --- | --- | --- |
| Lipid | 8.48e<sup>-3</sup> | 4.74e<sup>-3</sup> | 1.79 | 0.094 |
| Type | -5.89e<sup>-6</sup> | 3.80e<sup>-5</sup> | -0.155 | 0.879 |
| Lipid*Type | -3.68e<sup>-3</sup> | 5.25e<sup>-3</sup> | -0.701 | 0.494 |

**Table 6:**
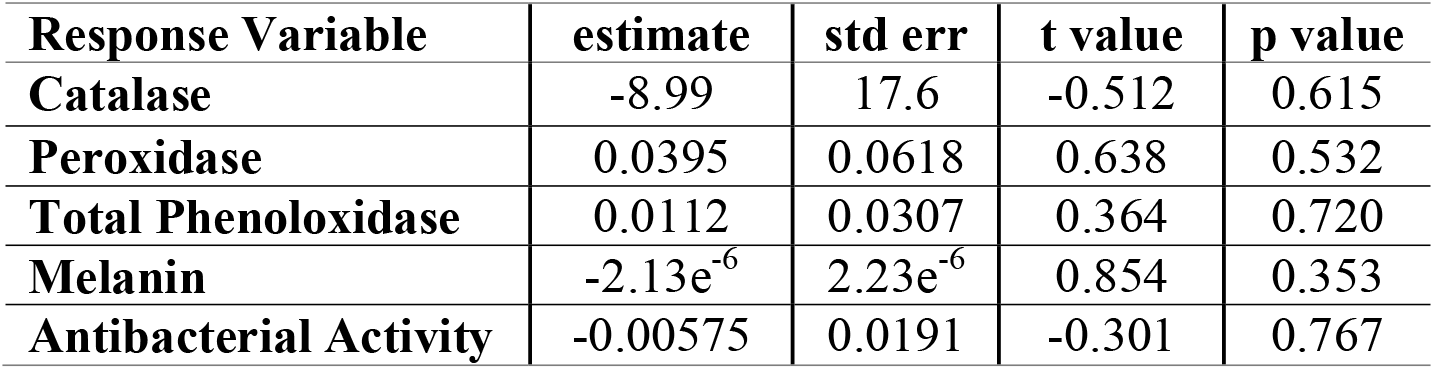
Results of linear model of the effects of sperm output on measured immune metrics.

## Discussion

Coral reefs globally are undergoing rapid, unprecedented declines due to anthropogenic climate change and associated disease outbreaks (Hoegh-Guldberg et al. 2007; Johnston et al. 2022; Precht et al. 2016). Corals exhibit both inter- and intra-specific variation in susceptibility to disease and infection, but the ecological factors that drive susceptibility, are poorly understood (Fuess et al. 2017; Mydlarz 2016). Few studies have investigated trade-offs between key life-history traits and immunity in cnidarians (Alvarez-Filip et al. 2022; Pinzon et al. 2014b; Schlecker et al. 2022; Weil et al. 2009). Here, we used the facultatively symbiotic coral *Astrangia poculata* to investigate potential reproductive-immune trade-offs in scleractinian corals, and the impacts of variation in symbiont density and energy budgets on these trade-offs.

Our results found no link between variation in symbiont density and any measured physiological trait (energetic budget, immune activity, reproductive output). Most notably, we observed no significant difference in total lipid and carbohydrate content as a result of variable symbiont density in adult coral colonies. While this is in agreement with a previous study of *A. poculata* (Szmant-Froelich & Pilson 1980), it is in opposition to conventional thinking regarding symbiosis, which would suggest that increased symbiont density results in increased energetic reserves (Changsut et al. 2022; Hughes et al. 2010; Pupier et al. 2019). We propose two alternative hypotheses to explain these patterns: first, it is possible corals with lower symbiont density compensate via increased heterotrophic feeding. Indeed, heterotrophic feeding has been demonstrated to increase physiological traits in other facultatively symbiotic corals (Aichelman et al. 2016). Alternatively, the total lipid and carbohydrate assays used here may not have fully captured host energetic budget. Future studies should investigate energetic reserves, in both the presence and absence of heterotrophy, and use additional methods to understand energetic budgets (i.e., metabolomics, carbon transfer, etc.).

Next, we examined links between energetic budget and other physiological traits, regardless of symbiont density. The only physiological trait associated significantly with energetic budget was melanin concentration. The observed association between melanin concentration and total lipids and carbohydrates is likely indicative of the high cost of melanin production (Gasparini et al. 2009; Talloen et al. 2004). Melanin production is the result of a complex pathway; beginning with pathogen recognition and continuing through a series of protein cascades leading to the production of melanin (Cerenius et al. 2010; Palmer et al. 2008). The complexity of the pathway is likely associated with high metabolic costs. To such end, previous studies have documented negative associations between lipid reserves and melanin concentration following immune stimulation (Sheridan et al. 2014). In contrast, the other immune metrics measured here are single molecules/compounds, and consequently likely require significantly less metabolic input (Traylor-Knowles & Connelly 2017). The low cost of investment in these components could explain the lack of strong correlation between our metrics of energetic budget and these immune parameters.

Finally, we examined potential tradeoffs between reproductive investment and immune activity. Contrary to our hypothesis, we observed no significant trade-offs between constitutive immunity and reproduction. We can posit several hypotheses to explain this observation. First, the timing of our experiment may not have properly reflected peak immune investment. Gametogenesis in *A. poculata* begins in early March-April and continues through June and July, with colonies spawning in early August/September (Szmant-Froelich et al. 1980). It is possible that allocation of resources to reproduction could have been higher at earlier points of gametogenesis, as opposed to at the time of sampling in August (Szmant-Froelich et al. 1980). Second, we chose here to measure sperm density due to sampling logistics. Sperm production is typically considered to be less energetically costly than production of eggs (Hayward & Gillooly 2011; Parker 1970; Parker 1982); it is possible that reproductive tradeoffs would be more evident when considering females. Finally, our study does not account for multiple spawning events, specifically spawning events that may have occurred naturally prior to sample collection. Some coral species spawn multiple times throughout a season, and exact timing of *Astrangia* spawning in natural environments is unknown (Szmant-Froelich et al. 1980). Thus, it is possible that the collected *Astrangia* colonies had previously spawned *in situ*, and the induced events were a secondary release with reduced sperm output not reflective of true reproductive investment. Congruent with this possibility, several of our colonies had negligible measured sperm output. Future studies combining histology with immunological assays at different points of gametogenesis in both male and female colonies may clarify presence and timing of potential trade-offs.

It is also possible that nuances in our study design did not allow us to observe trade-offs. It must be noted that trade-offs occur in a multi-dimensional trait space (Lochmiller & Deerenberg 2000; Stearns 1989), and our experimental analyses only captured two of these traits. It is highly likely that more complicated trade-offs may occur that were not captured by our study. Approaches that properly reflect the multi-dimensional trait space associated with resource allocation will be necessary to fully disentangle the relationship between immunity and other costly processes. Additionally, our study measured a limited number of immune metrics; it is possible that trade-offs exist between reproduction and immunity but involve other components of the immune system. The coral immune system is complex, involving different processes with differing relative costs (Colditz 2008; Ivanina et al. 2018; Lochmiller & Deerenberg 2000; Seppala & Leicht 2013). It is therefore reasonable to assume that trade-offs are not equivalent across components, as has previously been observed in other systems (Adamo et al. 2001; Albery et al. 2019; Gershman et al. 2010; Lochmiller & Deerenberg 2000). Future studies should incorporate more comprehensive metrics of immunity using methods such as gene expression or proteomics.

## Conclusions

In sum, our results fail to document notable effects of variation in symbiont density and energetic budget on coral physiology. Furthermore, we find no evidence of immune-associated trade-off. Still, our results are an important first step in broadening general understanding of resource allocation theory in cnidarians, which can be applied to other organisms facing rapid environmental changes. The information provided here provides an important preliminary framework for future studies of immunological trade-offs in marine invertebrates and the potential effects of variation in energetic budget on these patterns.

## Acknowledgements

The authors would like to thank Koty Sharp, Alicia Shickle, and members of the Roger Williams University Wet Lab for use of their facilities, assistance in collection of corals for this experiment, and execution of spawning/larval husbandry.

