## Supplementary figures and images for "Characterization of trade-offs between immunity and reproduction in *Astrangia poculata*"

### Fig 1

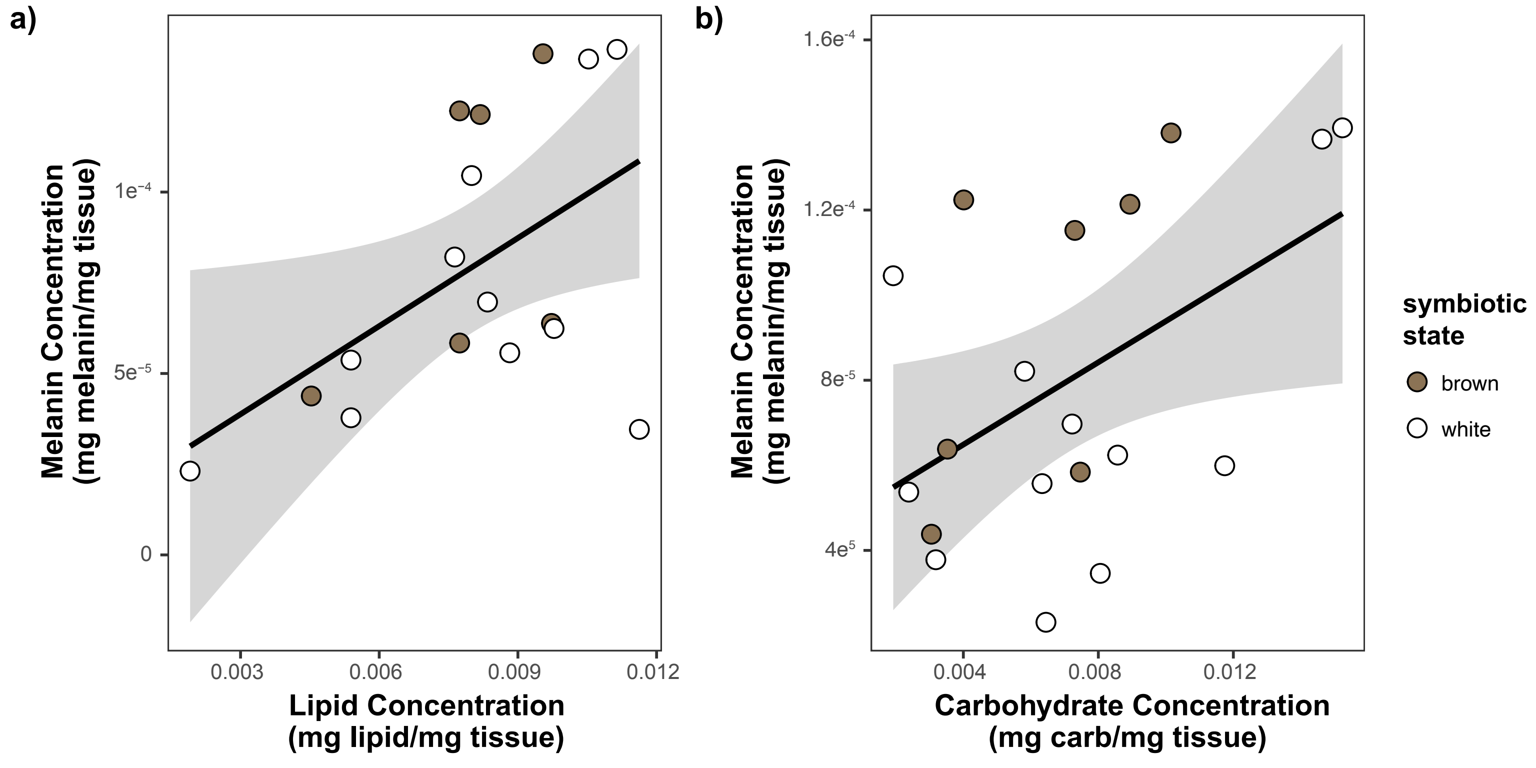
